# DEN-IM: Dengue Virus identification from shotgun and targeted metagenomics

**DOI:** 10.1101/628073

**Authors:** C I Mendes, E Lizarazo, M P Machado, D N Silva, A Tami, M Ramirez, N Couto, J W A Rossen, J A Carriço

## Abstract

Dengue virus (DENV) represents a public health and economic burden in affected countries.
The availability of genomic data is key to understand viral evolution and dynamics, supporting improved control strategies. Currently, the use of High Throughput Sequencing (HTS) technologies, which can be applied both directly to patient samples (shotgun metagenomics) and to PCR amplified viral sequences (targeted metagenomics), is the most informative approach to monitor the viral dissemination and genetic diversity.

Despite many advantages, these technologies require bioinformatics expertise and appropriate infrastructure for the analysis and interpretation of the resulting data. In addition, the many software solutions available can hamper reproducibility and comparison of results.

Here we present DEN-IM, a one-stop, user-friendly, containerised and reproducible workflow for the analysis of DENV sequencing data, both from shotgun and targeted metagenomics approaches. It is able to infer DENV coding sequence (CDS), identify serotype and genotype, and generate a phylogenetic tree. It can easily be run on any UNIX-like system, from local machines to high-performance computing clusters, performing a comprehensive analysis without the requirement of extensive bioinformatics expertise.

Using DEN-IM, we successfully analysed two DENV datasets. The first comprised 25 shotgun metagenomic sequencing samples of varying serotype and genotype, including a spiked sample containing the existing four serotypes. The second dataset consisted of 106 targeted metagenomics samples of DENV 3 genotype III where DEN-IM allowed detection of the intra-genotype diversity.

The DEN-IM workflow, parameters and execution configuration files, and documentation are freely available at https://github.com/B-UMMI/DEN-IM.

## 1 Key points

- Understanding DENV transmission in populations where the infection is endemic through the use of metagenomics is a most promising strategy to get insight into the dissemination of the virus and to support measures to prevent or decrease further spread.
- So far, the analysis, interpretation and dissemination of results encounters computational challenges, and requires appropriate expertise and infrastructure, which poses a challenge for the implementation of metagenomic approaches.
- We present DEN-IM, a reproducible, containerised and user-friendly workflow for the identification and characterisation of DENV from shotgun and targeted metagenomics data.
- The HTML reports obtained with DEN-IM can be easily shared, facilitating comparison of results from across the globe.

## 2 Background

The Dengue virus (DENV), a single-stranded positive-sense RNA virus belonging to the *Flavivirus* genus, is one of the most prevalent arboviruses and is mainly concentrated in tropical and subtropical regions. Infection with DENV results in symptoms ranging from mild fever to haemorrhagic fever and shock syndrome [1]. Transmission to humans occurs through the bite of Aedes mosquitoes namely *Aedes aegypti* and *Aedes albopictus* [2] In 2010, it was predicted that the burden of dengue disease reached 390 million cases per year worldwide [3]. The high morbidity and mortality of dengue makes it the arbovirus with the highest clinical significance [4].

The viral genome of ~11,000 nucleotides, consists of a Coding Sequence (CDS) of approximately 10.2 Kb that is translated into a single polyprotein encoding three structural proteins (capsid – C, premembrane – prM, envelope – E) and seven non-structural proteins (NS1, NS2A, NS2B, NS3, NS4A, NS4B and NS5. Additionally, the genome contains two Non-Coding Regions (NCRs) at their 5’ and 3’ ends [5].

DENV can be classified into four serotypes (1, 2, 3 and 4), differing from each other by 25% to 40% at the amino acid level. They are further classified into genotypes that vary by up to ~3% at the amino acid level [2]. The DENV-1 serotype comprises five genotypes (I-V), DENV-2 groups six (I-VI, also named American, Cosmopolitan, Asian-American, Asian II, Asian I and Sylvatic), DENV-3 four (I-III and V), and DENV-4 also four (I-IV).

DENV is a significant public health challenge in countries where the infection is endemic due to the high health and economic burden. Despite the emergence of novel therapies and ecological strategies to control the mosquito vector, there are still important knowledge gaps in the virus biology and its epidemiology [2].

The implementation of a surveillance system relying on High Throughput Sequencing (HTS) technologies allows the simultaneous identification and surveillance of DENV cases. Due to the high sensitivity of these technologies, previous studies showed that viral sequences can be directly obtained from patient sera using a shotgun metagenomics approach [6]. Alternatively, HTS can be used in a targeted metagenomics approach in which a PCR step is used to pre-amplify viral sequences before sequencing. In recent years, HTS has been successfully used as a tool for identification of DENV directly from clinical samples [6,7]. This also allows the rapid identification of the serotype and genotype important for disease management as the genotype may be associated with disease outcome [8].

Several initiatives aim to facilitate the identification of the DENV serotype and genotype from HTS data. The Genome Detective project (https://www.genomedetective.com/) offers an online Dengue Typing Tool (https://www.genomedetective.com/app/typingtool/dengue/) relying on BLAST and phylogenetic methods in order to identify the closest serotype and genotype, but it requires as input assembled genomes in FASTA format. Alternatively, the same project offers a Genome Detective Typing Tool (https://www.genomedetective.com/app/typingtool/virus/) identifying viruses present in a sample.

We developed DEN-IM as a ready-to-use, one-stop, reproducible bioinformatic analysis workflow for the processing and phylogenetic analysis of DENV using paired-end raw HTS data. DEN-IM is implemented in Nextflow [9], a workflow manager software that uses Docker (https://www.docker.com) containers with pre-installed software for all the workflow tools. The DEN-IM workflow, as well as parameters and documentation, are available at https://github.com/B-UMMI/DEN-IM.

## 3 The DEN-IM Workflow

DEN-IM is a user-friendly automated workflow allowing the analysis of shotgun or targeted metagenomics data for the identification, serotyping, genotyping, and phylogenetic analysis of DENV. It is implemented in Nextflow, a workflow management system that allows the effortless deployment and execution of complex distributed computational workflows.

The workflow is composed of five blocks: 1. Quality Control and Trimming, 2. Retrieval of DENV sequences, 3. Assembly, 4. *in silico* Typing, and 5. Phylogeny, described in more detail in Figure 1. DEN-IM accepts as input raw paired-end sequencing data (FASTQ files), and informs the user with an interactive HTML report with information on the quality control, mapping, assembly typing and phylogenetic analysis, as well as all the output files of the whole pipeline.

**Figure 1.**
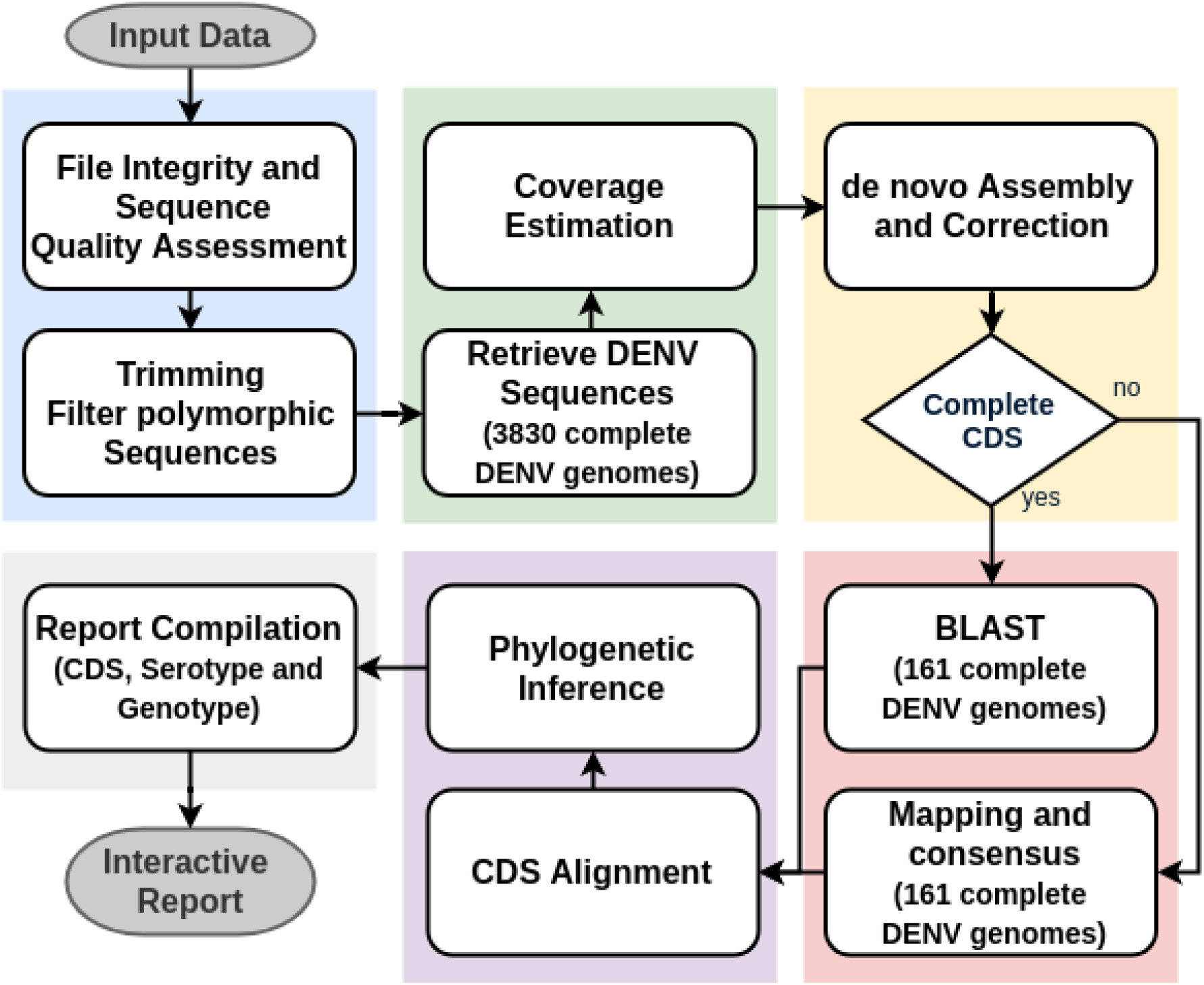
The DEN-IM workflow separated into five different components. The raw sequencing reads are provided as input to the first block (in blue), responsible for quality control and elimination of low-quality reads and sequences. After successful pre-processing of the reads, these enter the second block (green) for retrieval of the DENV reads using the mapping database of 3830 complete DENV genomes as reference. This block also provides an initial estimate of the sequencing depth. After the *de novo* assembly and assembly correction block (yellow), the coding sequences (CDSs) are retrieved and are then classified with the reduced complexity DENV typing database containing 161 sequences representing the known diversity of DENV serotypes and genotypes (red).If a complete CDS fails to be assembled, the reads are mapped against the DENV typing database and a consensus sequence is obtained for classification and phylogenetic inference. All CDSs are aligned and compared in a phylogenetic analysis (purple). Lastly, a report is compiled (gray) with the results of all the blocks of the workflow.

### 3.1 1. Quality Control and Trimming

The Quality Control (QC) and Trimming block starts with a process to verify the integrity of the input data. If the sequencing files are corrupted, the execution of the analysis of that sample is terminated.

The sequences are then processed by FastQC (https://www.bioinformatics.babraham.ac.uk/projects/fastqc/, version 0.11.7) to determine the quality of the individual base pairs of the raw data files. The low quality bases and adapter sequences are trimmed by Trimmomatic [10] (version 0.36). By default, Trimmomatic removes the Illumina Nextera, TruSeq2 and TruSeq3 adapter sequences that are detected, and relies on Phred scores [11] and FastQC analysis data to determine to which extend the 5’ and 3’ ends of the reads need to be trimmed. The crop and headcrop setting is calculated per sample, based on the GC ratio of each half of the read. By default, the window size is set to 5 nucleotides with a minimum average quality of 20. In addition, paired-end reads with a read length shorter than 55 nucleotides after trimming are removed from further analyses.

Lastly, the low complexity sequences, containing over 50% of poly-A, poly-N or poly-T nucleotides, are filtered out of the raw data using PrinSeq [12] (version 0.10.4).

### 3.2 2. Retrieval of DENV Sequences

In the second step, DENV sequences are selected from the sample using Bowtie2 [13] (version 2.2.9) and Samtools [14] (version 1.4.1). As a reference we provide the DENV mapping database, a curated DENV database composed of 3830 complete DENV genomes (see Methods, DENV Reference Database). A permissive approach is followed by allowing for mates to be kept in the sample even when only one read maps to the database in order to keep as many DENV derived reads as possible. The output of this step is a set of processed reads of putative DENV origin.

### 3.3 3. Assembly

DEN-IM applies a two assembler approach to generate assemblies of the DENV CDS. To obtain a high confidence assembly, the processed reads are first *de novo* assembled with SPAdes [15] (version 3.12.0). If the full CDS fails to be assembled into a single contig, the data is re-assembled with MEGAHIT assembler [16] (version 1.1.3), a more permissive assembler developed to retrieve longer sequences from metagenomics data. The resulting assemblies are corrected with Pilon [17] (version 1.22) after mapping the processed reads to the assemblies with Bowtie2 [13].

If more than one complete CDS is present in a sample, each of the sequences will follow the rest of the DEN-IM workflow independently. If no full CDS is assembled neither with SPAdes nor with MEGAHIT, the processed reads are passed on to the next step for consensus generation by mapping, effectively constituting DEN-IM’s two pronged approach using both assemblers and mapping.

### 3.4 4. Typing

For each DENV complete CDS, the serotype and genotype is determined with the Seq_Typing tool (https://github.com/B-UMMI/seq_typing, version 2.0) [18] using BLAST [19] and the custom Typing database of DENV containing 161 complete sequences (see Methods, DENV Reference Database). The BLAST results are first cleaned to get the best hit for each sequence in the database, based on alignment length, similarity, e-value and number of gaps. The tool determines which reference sequence is more closely related to the query based on the identity and length of the sequence covered, returning the serotype and genotype of the reference sequence for the purpose of classifying the query.

If a complete CDS fails to be obtained through the assembly process, the processed reads are mapped against the DENV typing database, with Bowtie2 [13], using Seq_Typing tool, with similar criteria for coverage and identity to those used with the BLAST approach. If a type is determined, the consensus sequence obtained follows through to the next step in the workflow. Otherwise, the sample is classified as Non-Typable and its process terminated.

### 3.5 5. Phylogeny

All DENV complete CDSs and consensus sequences analysed in a workflow execution are aligned with MAFFT [20] (version 7.402), in auto mode and with orientation being automatically determined and adjusted. With the resulting alignment, a Maximum Likelihood tree is inferred with RaXML [21] (version 8.2.11), using as default the GTR-Γ substitution model and a 500 times bootstrap.

Optionally, the closest reference sequence in the DENV typing database to each analysed sample can be retrieved and included in the phylogenetic analysis.

## 4 Workflow Execution

The DEN-IM workflow can be executed in any UNIX-based system, from local machines to high-performance computing clusters (HPC) facilities with Nextflow and a container engine installation, such as Docker (https://www.docker.com/, Shifter [22] or Singularity [23].

Due to its Nextflow implementation, DEN-IM allows for out-of-the-box high-level parallelization and offers direct support for distributed computational environments. DEN-IM integrates Docker containerised images for all the tools necessary for its execution, ensuring reproducibility and the tracking of both software code and version, regardless of the operating system used. The Docker images provided are also compatible with other container engines.

Users can customise the workflow execution either by using command line options or by modifying a simple plain-text configuration file (params.config). The version of each of the tools used in DEN-IM can be changed by providing new container tags in the appropriate configuration file (containers.config). The resources for each process can also be changed (resources.config). To make the execution of the workflow as simple as possible, a set of default parameters and directives is provided.

The local installation of the DEN-IM workflow, including the docker containers with all the tools needed and the curated DENV database, requires 15 Gigabytes of free disk space. The minimum requirements to execute the workflow are at least 5 Gigabytes of memory and 4 CPUs, although 7 Gigabytes of memory is advised. The disk space required for execution depends greatly on the size of the input data, but for the datasets used in this article DEN-IM generates approximately 20 Gb data per Gb of input data.

## 5 Output and Report

The output files of all tools are stored in the ‘results’ folder in the directory of the DEN-IM execution. The execution log file for each component, as well as a log file for the execution of the workflow, are also available.

DEN-IM creates an interactive HTML report (Figure S1), stored in the ‘pipeline_results’ directory, containing all the information in the results divided into four sections: report overview, tables, charts and phylogenetic tree. The report can be easily exchanged between collaborators by compressing and sharing the “pipeline_report” folder.

The report overview contains information about the number of samples in the analysis. It allows for selection, filtering and highlighting of particular samples and tools in the workflow.

The table section contains the results and statistics the quality control, assembly, read mapping and *in silico* typing results. The ***in silico* typing table** contains the results of the serotype and genotype of each CDS analysed, as well as identity and coverage and GenBank ID for the closest reference in the DENV typing database.

The **quality control table** shows information regarding the number of raw base pairs and number of reads in the raw input files and the percentage of trimmed reads. The **mapping table** includes the results for the mapping of the trimmed reads to the DENV mapping database, including the overall alignment rate, and an estimation of the sequence depth including only the DENV reads. For the **assembly statistics table**, the number of CDSs in each sample is included, and the number of contigs and the number of assembled base pairs generated by either SPAdes or MEGAHIT assemblers. The number of contigs and assembled base pairs after correction with Pilon is also presented in the table. Warning and fail messages are included in each table, as well as the ranking of the value in the cell in relation to other values in the same column, which are shown with a grey bar (Figure S1).

The **assembled contig size distribution scatter plot** is available in the chart section, showing the contig size distribution for the Pilon corrected assembled CDSs.

Lastly, a **phylogenetic tree** is included, rooted at midpoint for visualisation purposes, and with each tip coloured according to the genotyping results. If the option to retrieve the closest typing reference is selected, these sequences are also included in the tree with respective typing metadata. The tree can be displayed in several conformationsprovided by Phylocanvas JavaScript library (http://phylocanvas.net, version 2.8.1) and it is possible to zoom in or collapse selected branches. The support bootstrap values of the branches can be displayed and the tree can be exported as a Newick tree file or as a PNG image.

## 6 Results

To evaluate the DEN-IM workflow performance, we analysed two datasets, one containing shotgun metagenomics sequencing data of patient samples and another with targeted metagenomics sequencing data obtained from Parameswaran *et al* [24].

### 6.1 The Shotgun Metagenomics Dataset

We analysed 22 shotgun metagenomics paired-end short-read Illumina sequencing datasets of clinical positive dengue cases (Table 1)(identified as such by positive qRT-PCR/RT-PCR test and seroconversion test), positive control (purified from DENV culture) and one negative control (blank), and a spiked sample containing the 4 DENV serotypes.

**Table 1.**
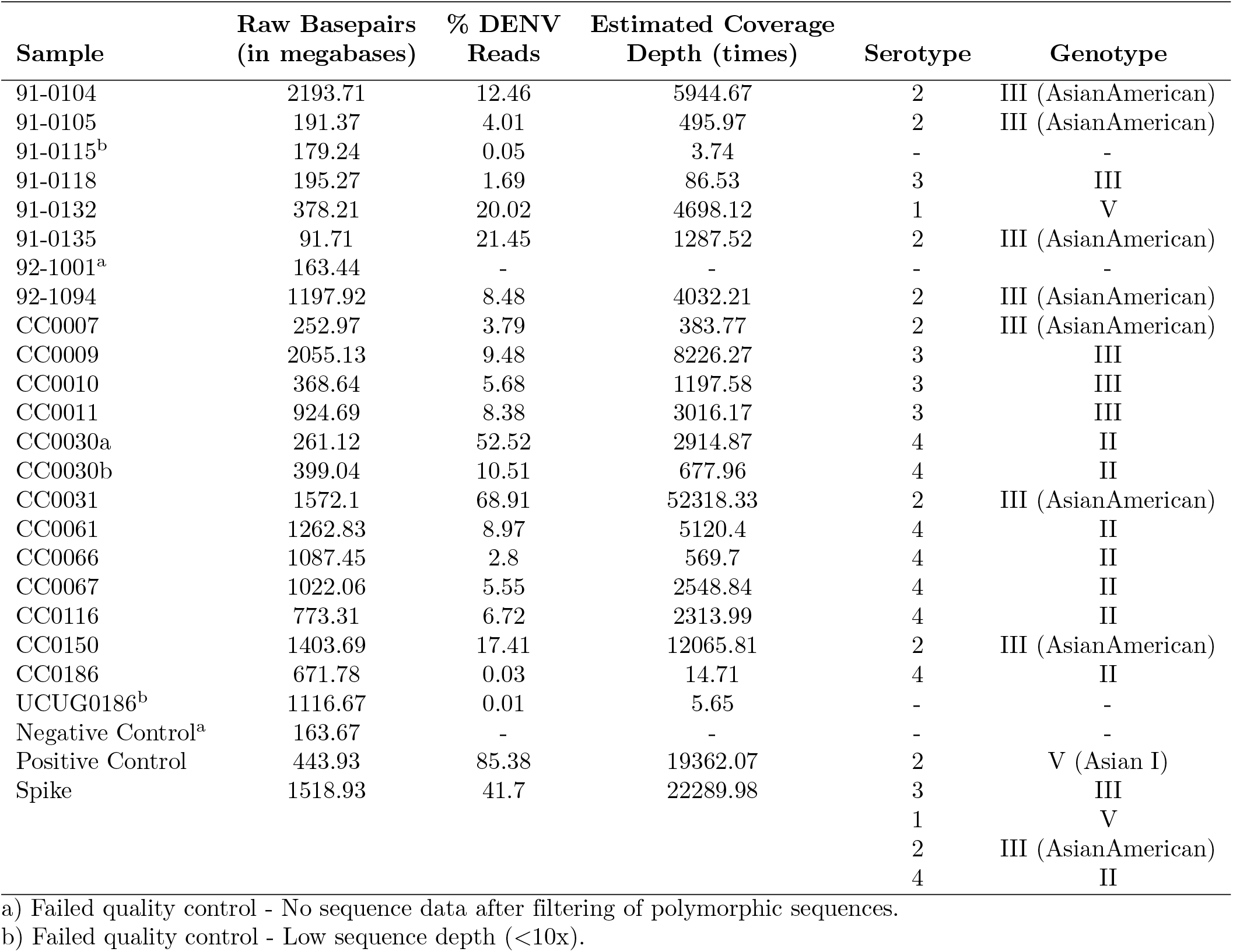
Number of raw base pairs, overall alignment rate against the DENV mapping database, estimated coverage depths and serotype and genotype for 25 shotgun metagenomics sequencing samples.

The workflow was executed using the default parameters and directives for resources, with the option to include the closest typing references in the final tree.

The negative control and the 92-1001 sample has no reads after trimming and filtering of low complexity reads, therefore they were removed from further analysis.

When mapping to the DENV mapping database, the percentage of DENV reads in the 21 clinical samples, positive control and spiked sample passing QC ranged from 0.01% (sample UCUG0186) to 85.38% (sample Positive Control). After coverage depth estimation, the analysis of the samples 91-0115 and UCUG0186 was terminated since they did not meet the threshold criterion of having an estimated depth of coverage of 10x.

In the assembly module, the remaining 19 clinical samples, the spiked sample and the positive control were assembled with DEN-IM’s two assembler approach. Twenty-four full CDS were assembled (Figure 2), despite originally having DENV reads content as low as 0.03% of the total number of reads. Sixteen samples, including the spiked sample and the positive control, were assembled in the first step with the SPAdes assembler, and five in the second with the MEGAHIT assembler. In the spiked sample, all four CDSs were successfully assembled and recovered.

**Figure 2.**
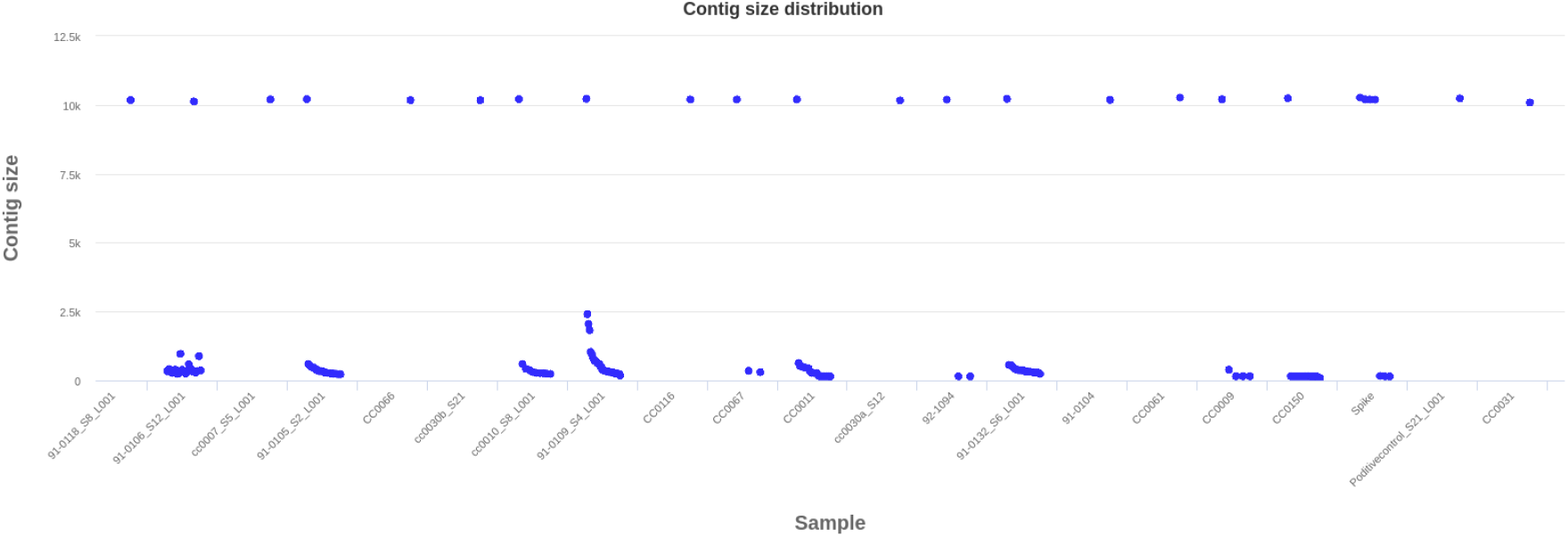
Contig size distribution for the shotgun metagenomics sequencing dataset. Each dot depicts an assembled DENV contig. Above the 10Kb are full CDS of DENV.

The serotype and genotype was successfully determined for the 24 DENV CDSs by BLAST (Figure S1, Table 1). The most common were serotype 2 genotype III (AsianAmerican) and serotype 4 genotype II, with 8 samples each (33.33%), followed by serotype 3 genotype III (n=5, 20.83%), serotype 1 genotype V (n=2, 8.33%) and serotype 2 genotype V (Asian I) (n=1, 4.17%). All CDSs recovered and the respective closest reference genome in the typing database were aligned and a maximum likelihood phylogenetic tree was obtained to visualise the relationship between the samples (Figure 3). There was a perfect concordance between the results of serotyping and genotyping and the major groups in the tree.

**Figure 3.**
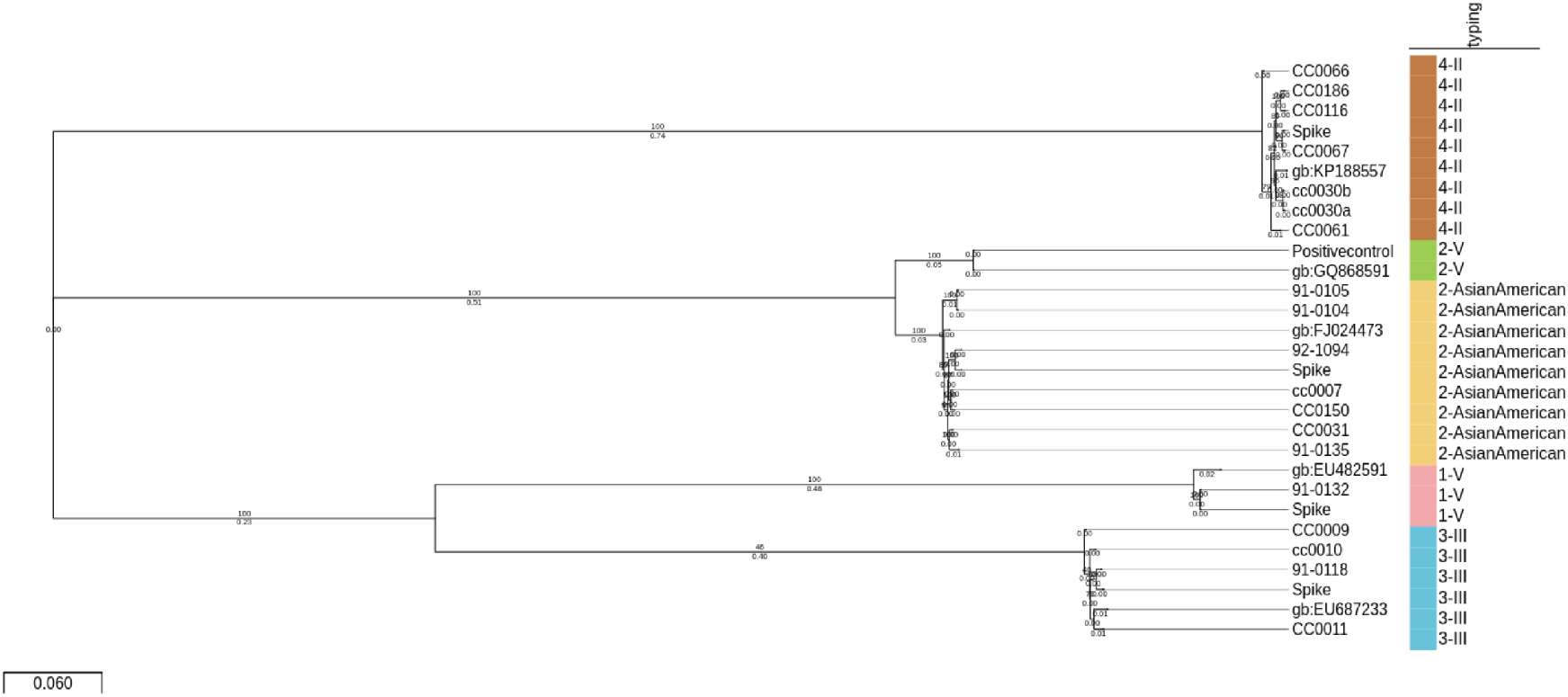
Maximum Likelihood tree in the DEN-IM report for the 24 complete CDSs (n=21 samples) obtained with the metagenomics dataset and the respective closest references in the typing database (identified by their GenBank ID). The tree is midpoint rooted for visualisation purposes. The colours depicts the DENV genotyping results.

### 6.2 The Targeted Metagenomics Dataset

To validate DEN-IM’s performance in a targeted metagenomics approach, 106 HTS datasets of PCR products using primers targeting DENV-3 [24] were analysed. As with the shotgun metagenomics dataset, the workflow was executed using the default parameters and directives for resources. This time not including the closest reference genome present in the typing database in the multiple sequence alignment due to the large number of input samples.

No samples failed the quality control block. The proportion of DENV reads ranged from 24.72% (SRR5821236) to 99.81% (SRR5821202) of the total processed reads. The samples with less than 70% DENV DNA were taxonomic profiled with the Kraken2 [25] with the minikraken2_v2 database (ftp://ftp.ccb.jhu.edu/pub/data/kraken2_dbs/minikraken2_v2_8GB_201904_UPDATE.tgz) and the source of the contamination was determined to have come largely from Human DNA (Table S2).

Of the 106 samples, 43 (40.60%) managed to assemble a complete CDS sequence (Table S1) whereas a mapping approach was used for the remaining 63 samples (59.90%) and a consensus CDS was generated. For the assembled CDSs, all but one were assembled with MEGAHIT after not producing a full CDS with SPAdes. Moreover, pronounced variation on the size of the assembled contigs is evident in the contig size distribution plot (Figure 4).

**Figure 4.**
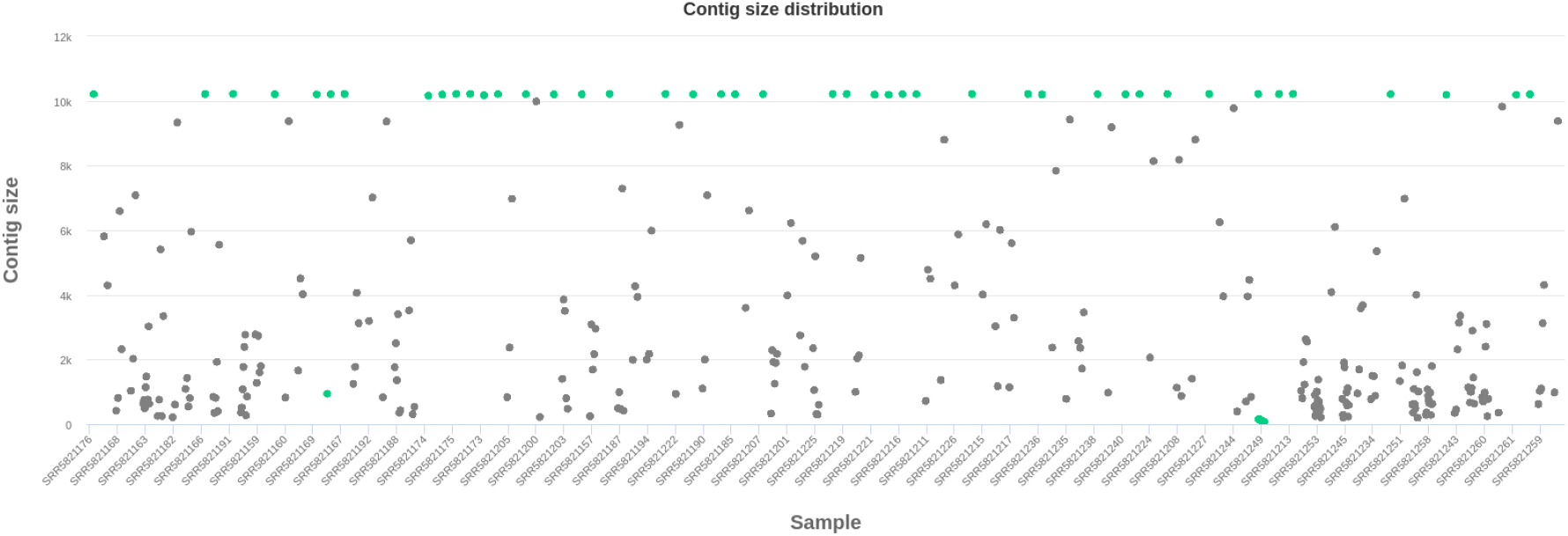
Contig size distribution of the targeted metagenomics dataset. Each dot depicts an assembled DENV contig. Above the 10Kb are full CDS of DENV. Contigs belonging from samples that assembled a complete DENV CDS are highlighted in green, whereas the remaining are coloured in grey.

All 106 CDSs recovered belonged to serotype 3 genotype III. Despite the same classification, the maximum likelihood tree indicates that there is detectable genetic diversity within the samples (Figure 5).

**Figure 5.**
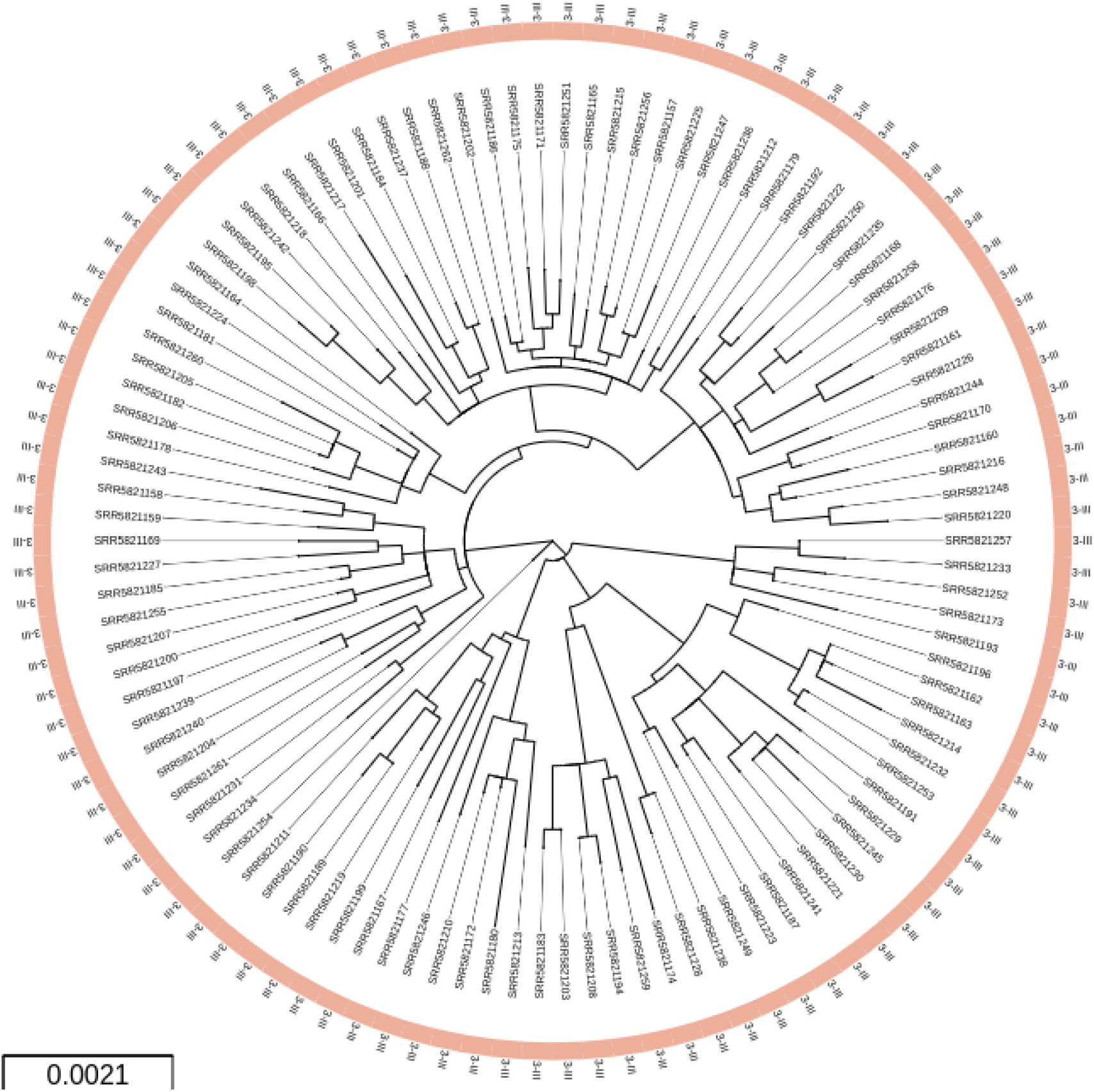
Maximum likelihood circular tree in the DEN-IM report for the 106 complete CDSs obtained with the targeted metagenomics dataset (n=106). All samples belong to serotype 3 genotype III.

## 7 Discussion

We have successfully analysed two DENV datasets, one comprising 25 shotgun metagenomics sequencing data and 106 targeted metagenomics data.

In the first dataset, we recovered 24 CDSs from 19 clinical samples, a spiked sample and a positive control that were correctly serotyped and genotyped. Besides the negative control, 3 samples did not return typing information due to failing quality checks. In one case (92-1001), no DENV reads after quality control processing were detected as all the reads contained highly repetitive sequences (AAA; TTTT) and were filtered out. The two others (91-0115 and UCUG0186) had a low proportion of DENV reads (0.05% and 0.01%) and an estimated depth of coverage lower than the 10x threshold criterion (3.17x and 5.65x, respectively). Sequence data of sample CC0186 contained only 960 DENV reads (0.03%), but these were successfully assembled into a CDS with an estimated depth of coverage of 14.71x.

The proportion of DENV reads was the metagenomic samples is very variable. This may reflect the viral load in patients in which DENV was detected by PCR. In the spiked sample, containing 4 distinct DENV serotypes, all four were correctly detected despite not being present in equal concentrations(see Methods, Targeted Metagenomics Sequencing Data). This resulted in different coverage of each serotype CDS (2032.31 times coverage for DENV-2, 229.02 times coverage for DENV-1, 76.47 times coverage for DENV-3 and 29.78 times coverage for DENV-4), in accordance with the ranking order of RT-PCR results. It highlights the potential of the DEN-IM workflow to accurately detect and recover multiple DENV genomes from samples with DENV co-infection, even if the serotypes are present in low abundance. Indeed, recent studies from areas of high endemicity suggest that co-infection with multiple DENV serotypes may frequently occur [26] [27] and the co-circulation of different DENV strains of the same serotype, but distinct genotypes, in these areas [26] raises the possibility of simultaneous infection with more than one genotype.

When analysing the targeted metagenomics dataset, only 43 CDS out of 106 samples were *de novo* assembled. For the remaining 63 samples, consensus sequences were obtained through mapping. In all samples DENV 3-III was correctly identified, demonstrating the success of DEN-IM’s two pronged approach of combining assemblers and mapping. We suggest that the lower assembly success of the targeted metagenomics data may be related to errors during the amplification process resulting in low quality reads ends which are then trimmed by the quality control block potentially affecting the assembly process as the overlapping regions are diminished.

DEN-IM is built with modularity and containerisation as keystones, leveraging the parallelization of processes and guaranteeing reproducible analyses across platforms. The modular design allows for new modules to be easily added and tools that become outdated can be easily updated, ensuring DEN-IM’s sustainability. The software versions are described in the Nextflow script and configuration files, and in the dockerfiles for each container, allowing traceability of each step of data processing.

Being developed in Nextflow, DEN-IM runs on any UNIX-like system and provides out-of-the-box support for several job schedulers (e.g., PBS, SGE, SLURM) and integration with containerised software like Docker or Singularity. While it has been developed to be ready to use by non-experts, not requiring any software installation or parameter tuning, it can still be easily customised through the configuration files.

The interactive HTML reports (Figure S1) provide an intuitive platform for data exploration, allowing the user to highlight specific samples, filter and re-order the data tables, and export the plots as needed. Together with the workflow and software containers, a database containing 3830 complete DENV genomes for DENV sequence retrieval and a subset database with 161 curated DENV genomes for serotyping and genotyping are provided. While constructing these databases, the obstacles reported by Cuypers *et al* [28] were apparent, namely the lack of formal definition of a DENV genotype and the lack of a standardised classification procedure that could assign sequences to a previously defined genotypic/sub-genotypic clade [28]. Discrepancies between the phylogenetic relationship and the genotype assignment were frequent and, throughout this study, the classification of some strains within the ViPR database [29] was updated.

As suggested previously [28], further evaluation of the DENV classification will benefit future research and investigation into the population dynamics of this virus. Our typing approach was designed to use the currently accepted DENV classification. However, DEN-IM can be easily modified if a new DENV classification system is to be established in the future.

In conclusion, we provide a user-friendly workflow that makes it possible to analyse paired-end raw sequencing data from shotgun or targeted metagenomics for the presence, typing and phylogenetic analysis of DENV. The use of containerised workflows, together with shareable reports, will allow an easier comparison of results globally, promoting collaborations that can benefit the populations where DENV is endemic. The DEN-IM source code is freely available in the DEN-IM GitHub repository (https://github.com/B-UMMI/DEN-IM), which includes a wiki with full documentation and easy to follow instructions.

## 8 Potential implications

The burden of DENV disease is already large, but is still increasing as the risk of exposure to the virus is increasing, not only through travel to endemic areas but also due to the expansion of the geographic areas of the mosquito vectors and the disease [30].

The decreasing costs and wider availability of HTS makes it an ideal technology to monitor DENV transmission, including the direct processing of patient samples, either through the use of a more affordable targeted metagenomics approach or through shotgun metagenomics. Either way, a ready to use bioinformatics workflow that enables the reproducible analysis of DENV is particularly relevant.

DEN-IM was designed to perform a comprehensive analysis without the requirement of extensive bioinformatics expertise in order to generate either assemblies or consensus of full DENV CDSs and to identify the serotype and genotype of the DENV present in the sample. Although we did not exhaustively test the capacity of DEN-IM to detect co-infection with multiple DENV serotypes and genotypes, all four genotypes present in the spiked sample were accurately detected. This raises the possibility that DEN-IM can play a role in the identification of these cases whose prevalence is increasingly appreciated in highly endemic areas. Moreover, although being ready-to-use, the DEN-IM workflow can be easily customised to optimise the data analysis.

DEN-IM enables reproducible and collaborative research, benefiting a wide group of researchers regardless of their computational expertise and resources available.

## 9 Methods

### 9.1 DENV Reference Database

We have compiled a database of 3830 complete DENV genomes obtained from the NIAID Virus Pathogen Database and Analysis Resource (ViPR) in January 2019 [29] (http://www.viprbrc.org/). The sequences were distributed unevenly throughout the four DENV serotypes, with DENV-1 being the most represented with 1636 sequences (42.72%), followed by DENV-2 with 1067 sequences (27.86%), DENV-3 with 807 sequences (21.07%), and DENV-4 with 320 sequences (8.36%). The selection criteria for the search were as follows: a) complete genome sequence only, b) human host only, c) collection year (1950-2018). Data available from all countries was included and duplicated sequences were removed and only the sequences with sub-type data were kept. A representative of DENV serotype 1 genotype III was introduced (EF457905, recovered from monkey) as no representatives were available with the search criteria used. This genotype is Sylvatic and considered extinct [31] [32]. Additionally, any sample with IUPAC codes in the sequence provided were excluded.

In order to recover the maximum number of DENV reads from the input HTS data in the first mapping step (Figure 1), we maintained the database with the 3830 complete DENV genomes to retain as much diversity as possible. This database is referred as “DENV mapping database”.

For typing purposes, overly similar sequences in the collection were removed from the database by clustering the sequences in each serotype at 98% nucleotide similarity with CD-HIT [33], leaving 161 representative sequences of all described DENV serotypes and genotypes, with 46 DENV-1 sequences (Table S4), 63 DENV-2 (Table S5), 25 DENV-3 (Table S6) and 27 DENV-4 (Table S7). This database is referred as “DENV typing database”. This step is necessary to speed up the classification step for genotyping.

Phylogenetic analysis of typing collection was performed by aligning the reference genomes with MAFFT [20], in auto mode and with automatic sequence orientation adjustment. A phylogenetic tree was inferred with RAxML (version 8.12.11) [21] using the GTR-Γ substitution model and 500 times bootstrap. The resulting trees are available as supplemental material (Figures S2 to S5).

The sequence JF459993 from in the DENV-1 collection, as of April 2019, was annotated in ViPR as belonging to genotype IV, but in our analysis it clustered within genotype I clade (Figure S2). The classification of DENV-1 I was also obtained from GenomeDetective Dengue Typing Tool (https://www.genomedetective.com/app/typingtool/dengue/), so we proceeded to alter the annotation of this particular sample (Table S4). In order to harmonise dengue nomenclature, the system adopted uses Roman-numeric labels to identify the genotype, with the exception of Serotype 2 (Table S5), which used both Roman-numeric and geographic origin due to the widespread adoption of the latter.

### 9.2 Workflow Parameters

The short-read paired-end data is passed as input through the “*–fastq*” parameter, that by default is set to match all files in the “fastq” folder that match the pattern “*\*_R1,2\**”. In the process to verify the integrity of the paired-end raw sequencing data, the integrity of the input files is assessed by attempting to decompress and read the files. An estimation of the depth of coverage is also performed. By default, the input size (“*–genomeSize*”) is set to 0.012 Mb and the minimum coverage depth (“*–minCoverage*”) is set to 10. If any input file is found to be corrupt, its progression in the workflow is aborted.

In the FastQC and Trimmomatic module, FastQC (https://www.bioinformatics.babraham.ac.uk/projects/fastqc/) is run with the parameters “*–extract –nogroup –format fastq*”. FastQC will inform Trimmomatic [10] on how many bases to trim from the 3’and 5’ ends of the raw reads. By default, Trimmomatic uses the default set of Illumina adapters provided with the workflow but this behaviour can be overwritten with the “*–adapters*” parameter. The additional Trimmomatic parameters “*–trimSlidingWindow*”, “*–trimLeading*”, “*–trimTrailing*” and “*–trimMinLength*” can all be set to different values.

The removal of low complexity sequences is done with PrinSeq [12] using a custom parameter (“*–pattern*”), which by default is set to the value “A 50%; T 50%; N 50%”, removing sequences whose content is at least half composed of a polymeric sequence (A, T or N).

To retrieve the reads that map to the DENV reference database, Bowtie2 [13] is run with default parameters with the DENV mapping database as a reference. The reads and their mates that map to the reference are retrieved with “*samtools view -buh -F 12*” and “*samtools fastq*” commands. The DENV mapping database can be altered with the “*–reference*” parameter, or alternatively, a Bowtie2 index can be provided with the “*–index*” parameter. This allows for the workflow to work with other databases obtained through public and owned DENV genomes. The coverage estimation step is performed on the retrieved DENV reads with the same parameters are the first estimation (“—genomeSize=0.012” and “—minCoverage=10”).

In the assembly process, the retrieved DENV reads are firstly assembled with SPAdes Genome Assembler [15] with the options “*–careful –only-assembler –cov-cutoff* “. The coverage cutoff if dictated by the “*–spadesMinCoverage*” and “*–spadesMinKmerCoverage*” parameters, set to 2 by default. If the assembly with SPAdes fails to produce a contig equal or greater than the value defined in the “*–minimumContigSize*” parameter (default of 10000), the data is re-assembled with the MEGAHIT assembler [16] with default parameters. By default the k-mers to be used in the assembly in both tools (“*–spadesKmers*” and “*–megahitKmers*”) are automatically determined depending on the read size. If the maximum read length is equal or greater than 175 nucleotides, the assembly is done with the k-mers “55, 77, 99, 113, 127”, otherwise the k-mers “21, 33, 55, 67, 77” are used.

To correct the assemblies produced, the Pilon tool [17] is run after mapping the QC’ed reads back to the assembly with Bowtie2 and “*samtools sort*”. This process also verifies the coverage and the number of contigs produced in the assembly. The behaviour can be altered with the parameters “*–minAssemblyCoverage*”, “*–AMaxContigs*” and “*–genomeSize*”, set to “auto”, 1000 and 0.01 Mb by default. The first parameter, when set to ‘auto’, the minimum assembly coverage for each contig required is set to the 1/3 of the assembly mean coverage or to a minimum of 10x. The ratio of contig number per genome MB is calculated based on the genome size estimation for the samples.

The contigs larger than the value defined in the “*–size*” parameter (default of 10000 nucleotides) are considered to be complete CDSs and follow the rest to the workflow independently. If no complete CDS is recovered, the QC’ed read data is passed to the mapping to module that does the DENV typing database and consensus generation.

The serotyping and genotyping is performed with the Seq_Typing tool [18] with the command “*seq_typing.py assembly*”or “*seq_typing.py reads*”, using as reference the provided curated DENV typing database. It is possible to retrieve the genomes of the closest references and include them in the downstream analysis by changing the “–get_reference” option to “true”. By default this is not included in the analysis.

The CDSs, and the reference sequences if requested, are aligned with the MAFFT tool [20] with the options “*–adjustdirection –auto*” and a maximum likelihood phylogenetic tree is obtained with the RaXML tool [21] with the options “*-p 12345 -f -a*”. Additionally and by default, the substitution model (“*–substitutionModel*”) is set to “GTRGAMMA”, the bootstrap is set to 500 (“*–bootstrap*”) and the seed to “12345” (“*–seedNumber*”).

### 9.3 Shotgun Metagenomics Sequencing Data

Samples of plasma (n=9) and serum samples (n=13) from confirmed dengue symptomatic patients were collected in Venezuela between 2010-2015 (Table S3) (see Availability of supporting data and materials). DENV positivity was confirmed by either RT-qPCR [34] or nested RT-PCR [35].

As a positive control sample, the supernatant of a viral culture containing DENV-2 strain 16681 was used. The negative control sample consisted of DNA- and RNA-free water (Sigma-Aldrich, St. Louis, MO, USA).

A spiked sample was produced consisting of a mixture of four 5 *μ*l of cDNA isolated from clinical samples including all DENV serotypes (DENV-1 to -4). The viral cDNA for these samples was not in equal concentration and the viral copy number in the clinical samples was assessed by RT-PCR [35]. The results were as follow: DENV-2 with 1070000 copies/*μ*l, DENV-1 with 117830 copies/*μ*l, DENV-3 with 44300 copies/*μ*l and DENV-4 with 6600 copies/*μ*l.

The cDNA libraries were generated using either the NEBNext^®^ RNA First and Second strand modules and the Nextera XT DNA library preparation kit (NXT), or the TruSeq RNA V2 library preparation kit (TS). The libraries were sequenced in MiSeq and NextSeq instruments using 300-cycles v2 paired-end cartridges.

The DEN-IM workflow was executed with the raw sequencing data using the default parameters and resources in an HPC cluster with 300 Cores/600 Threads of Processing Power and 3 TB RAM divided through 15 computational nodes, 9 with 254 GB Ram and 6 with 126GB RAM.

### 9.4 Targeted Metagenomics Sequencing Data

The accession numbers for the 106 DENV-3 amplicon sequencing paired-end short-read datasets are available under BioProject PRJNA394021. The list of Run Accession IDs were obtained with NCBI’s RunSelector and the raw data was downloaded with the GetSeqENA tool (https://github.com/B-UMMI/getSeqENA).

The DEN-IM workflow was executed with the raw sequencing data with default parameters and resources in the same HPC cluster as the shotgun metagenomics dataset.

## 10 Availability of source code and requirements

Lists the following:

- Project name: DEN-IM
- Project home page: https://github.com/B-UMMI/DEN-IM
- Operating system(s): UNIX-like systems.
- Programming language: Nextflow, Python, Bash
- Other requirements: Java version 8 or highest. Docker/Singularity/Shifter
- License: GNU GPL v3
- Documentation and tutorials: https://github.com/B-UMMI/DEN-IM/wiki

## 11 Availability of supporting data and materials

The 106 DENV-3 targeted metagenomics sequencing paired-end short-read datasets are available under BioProject PRJNA394021. The 25 shotgun metagenomics dataset is available under Bioproject PRJNA474413 The accession number for all the samples in the shotgun metagenomics dataset are available in the supplemental material (Table S3).

## 12 Declarations

### 12.1 List of abbreviations

DENV: Dengue Virus
CDS: Coding Sequence
ORF: Open Reading Frame
NCR: Non-Coding Region
HPC: High-Performance Computing
HTS: High Throughput Sequencing
QC: Quality Control

### 12.2 Ethical Approval (optional)

This study followed international standards for the ethical conduct of research involving human subjects. Data and sample collection was carried out within the DENVEN and IDAMS (International Research Consortium on Dengue Risk Assessment, Management and Surveillance) projects. The study was approved by the Ethics Review Committee of the Biomedical Research Institute, Carabobo University (Aval Bioetico #CBIIB(UC)-014 and CBIIB-(UC)-2013-1), Maracay, Venezuela; the Ethics, Bioethics and Biodiversity Committee (CEBioBio) of the National Foundation for Science, Technology and Innovation (FONACIT) of the Ministry of Science, Technology and Innovation, Caracas, Venezuela; the regional Health authorities of Aragua state (CORPOSALUD Aragua) and Carabobo State (INSALUD); and by the Ethics Committee of the Medical Faculty of Heidelberg University and the Oxford University Tropical Research Ethics Committee.

### 12.3 Consent for publication

Not applicable.

### 12.4 Competing Interests

The authors declare that they have no competing interests.

### 12.5 Funding

C.I.M. was supported by the Fundação para a Ciência e Tecnologia (grant SFRH/BD/129483/2017). Erley Lizarazo received the Abel Tasman Talent Program grant from the UMCG, University of Groningen, Groningen, The Netherlands. This work was partly supported by the ONEIDA project (LISBOA-01-0145-FEDER-016417) co-funded by FEEI-Fundos Europeus Estruturais e de Investimento from Programa Operacional Regional Lisboa 2020 and by national funds from FCT-Fundação para a Ciência e a Tecnologia and UID/BIM/50005/2019, and by UID/BIM/50005/2019, project funded by Fundação para a Ciência e a Tecnologia (FCT)/ Ministério da Ciência, Tecnologia e Ensino Superior (MCTES) through Fundos do Orçamento de Estado.

### 12.6 Author’s Contributions

C.I.M., E.L., N.C., M.R., J.A.C. and J.W.A.R. designed the workflow. C.I.M implemented and optimised the workflow, created the Docker containers, and wrote the manuscript. M.P.M. implemented the DENV genotyping module in the workflow and D.N.S. contributed to the development of DEN-IM’s HTML report. E.L., A. T., and N.C. provided the shotgun metagenomics data used to test and validate the workflow and wrote the manuscript. A.T., N.C., M.R., J.A.C. and J.W.A.R. critically revised the article. All authors read, commented on, and approved the final manuscript.

## 13 Acknowledgements

The authors would like to thank Tiago F. Jesus and Bruno Ribeiro-Goncalves for their invaluable help with the Nextflow implementation. We would also like to thank Erwin C. Raangs from the UMCG for his assistance in the sequencing of the shotgun metagenomics dataset. Additionally, the authors thank Lize Cuypers, Krystof Theys, Pieter Libin and Gilberto Santiago for their assistance in DENV nomenclature and classification. This work was done in collaboration with the ESCMID Study Group on Molecular and Genomic Diagnostics (ESGMD), Basel, Switzerland.

---

**Figure S1.**
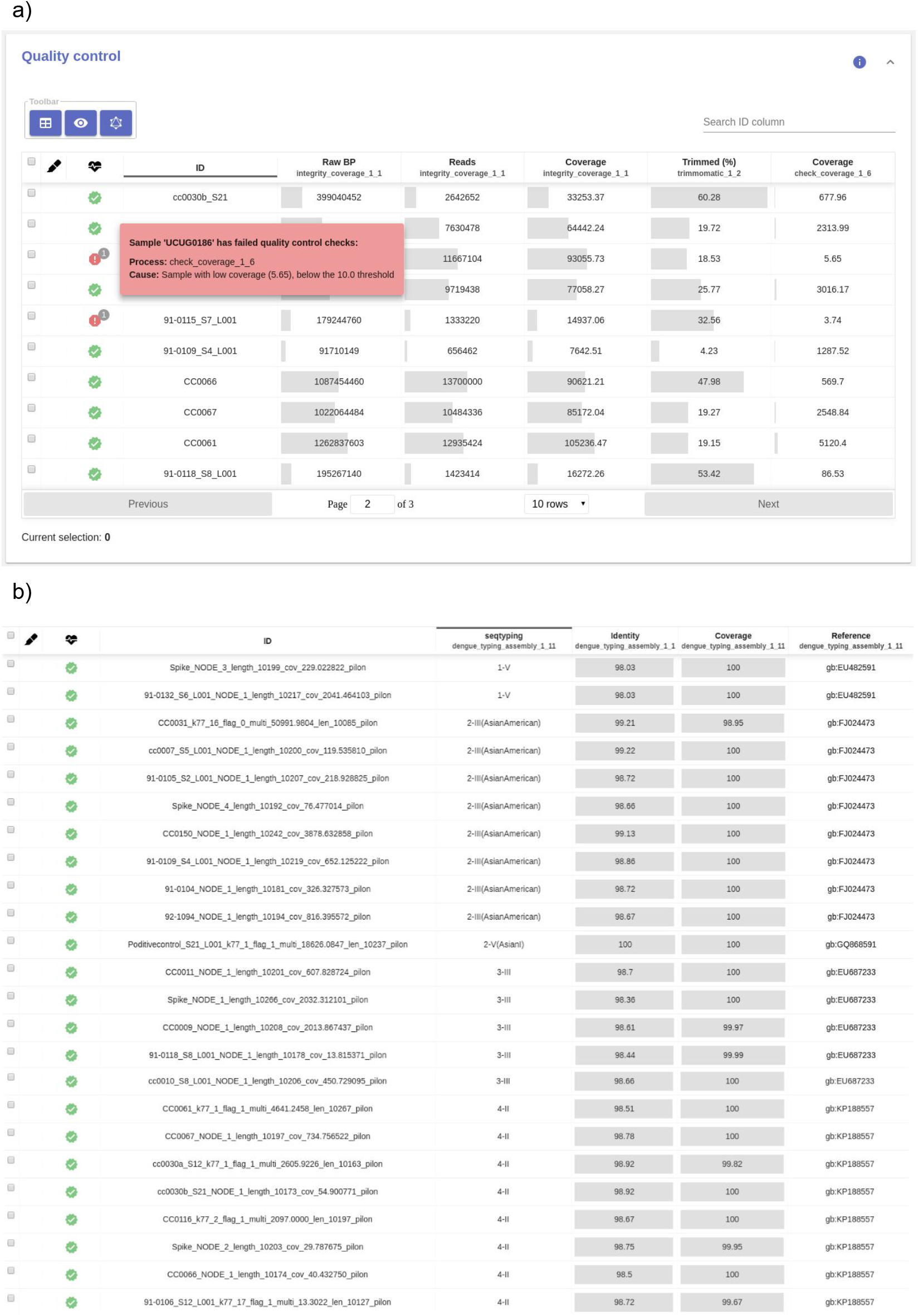
a) DEN-IM’s quality control report containing information of the number of basepairs and the number of reads for the analysed samples, the estimated coverage depth before and after mapping, and the percentage of reads in the input data that were trimmed. b)DEN-IM’s typing report for 24 CDSs recovered from the metagenomic dataset. The ID contains the CDS contig name, the typing result for serotype-genotype, the values for identity and coverage, and the GenBank ID of the closest reference in the Typing Database containing 161 complete DENV genomes.

**Figure S2.**
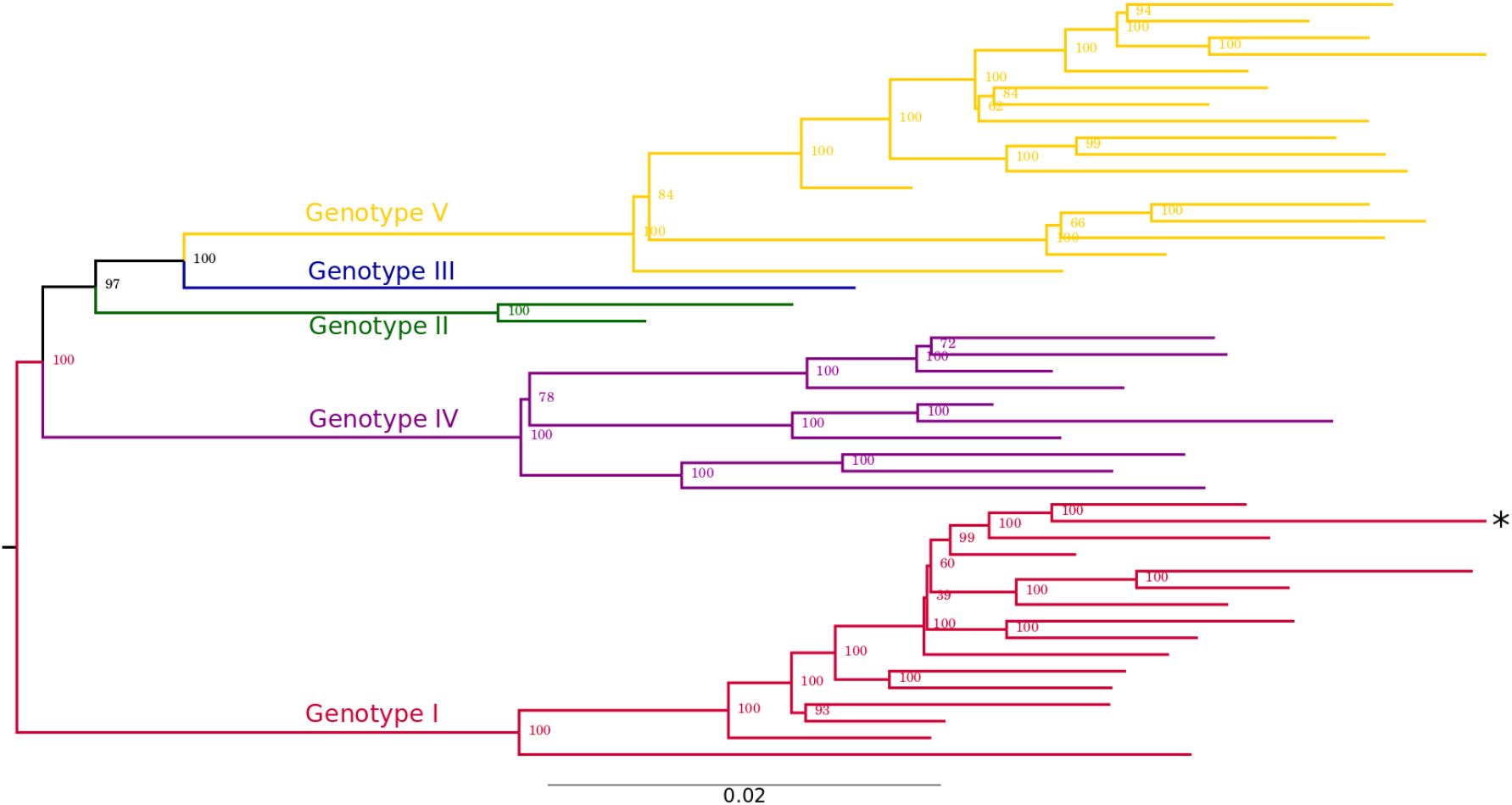
Maximum Likelihood inference of the multiple sequence alignment of the 46 DENV-1 complete genomes in the typing dataset. 1635 complete DENV-1 genomes were clustered at 98% nucleotide identity and the representative genomes were aligned with mafft. A maximum likelihood tree was infered with RAxML. The tree is coloured according to genotype (red: genotype I; green: genotype II; blue: genotype III; purple: genotype IV). The sample JF459993, marked with a star, is currently annotated in ViPR as belonging to genotype IV but, given to the good phylogenetic support, it was re-classified as belonging to the genotype I.

**Figure S3.**
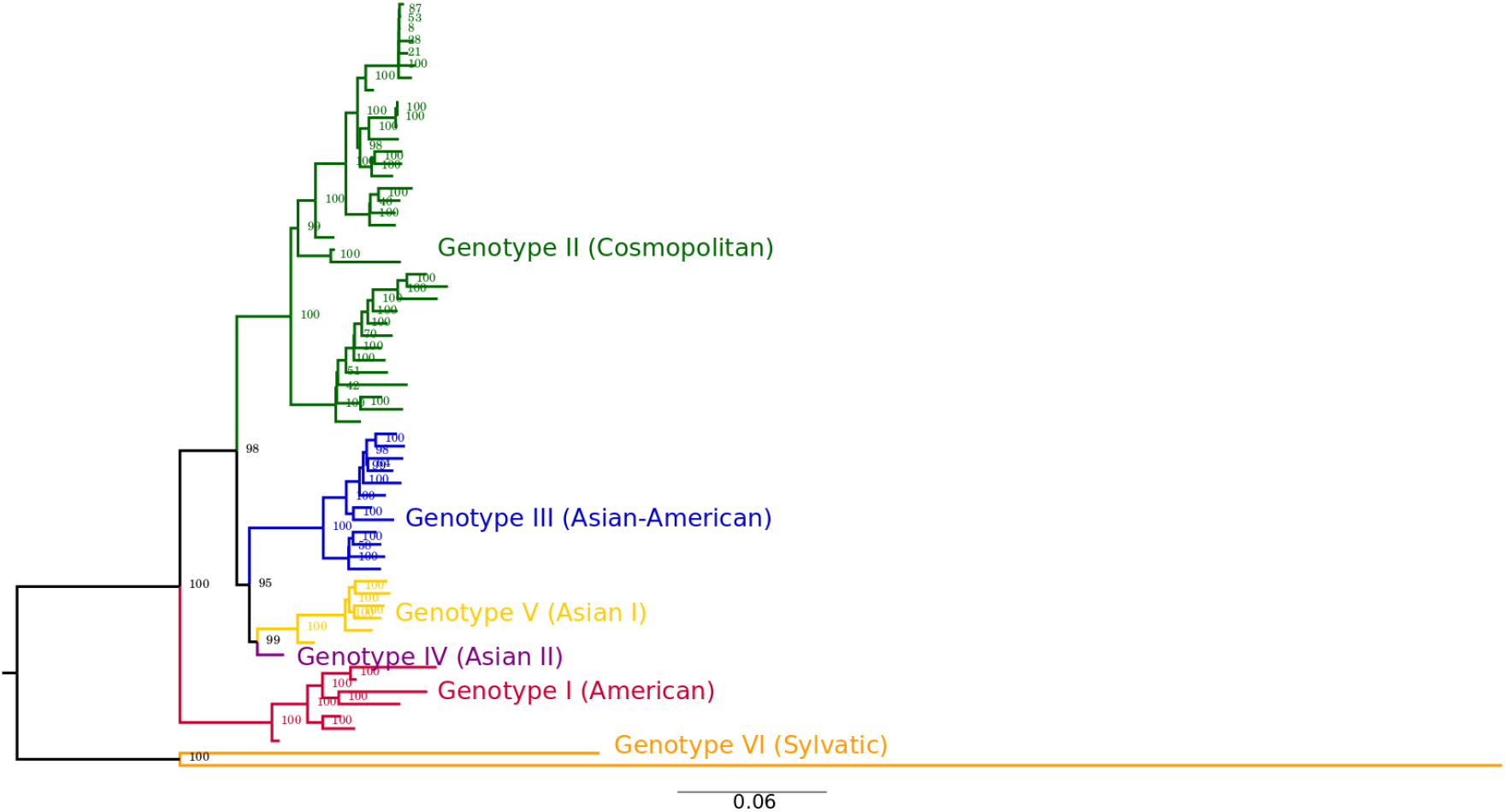
Maximum Likelihood inference of the multiple sequence alignment of the 63 DENV-2 complete genomes in the typing dataset. 1067 complete DENV-1 genomes were clustered at 98% nucleotide identity and the representative genomes were aligned with mafft. A maximum likelihood tree was infered with RAxML. The tree is coloured according to genotype (red: genotype I; green: genotype II; blue: genotype III; purple: genotype IV).

**Figure S4.**
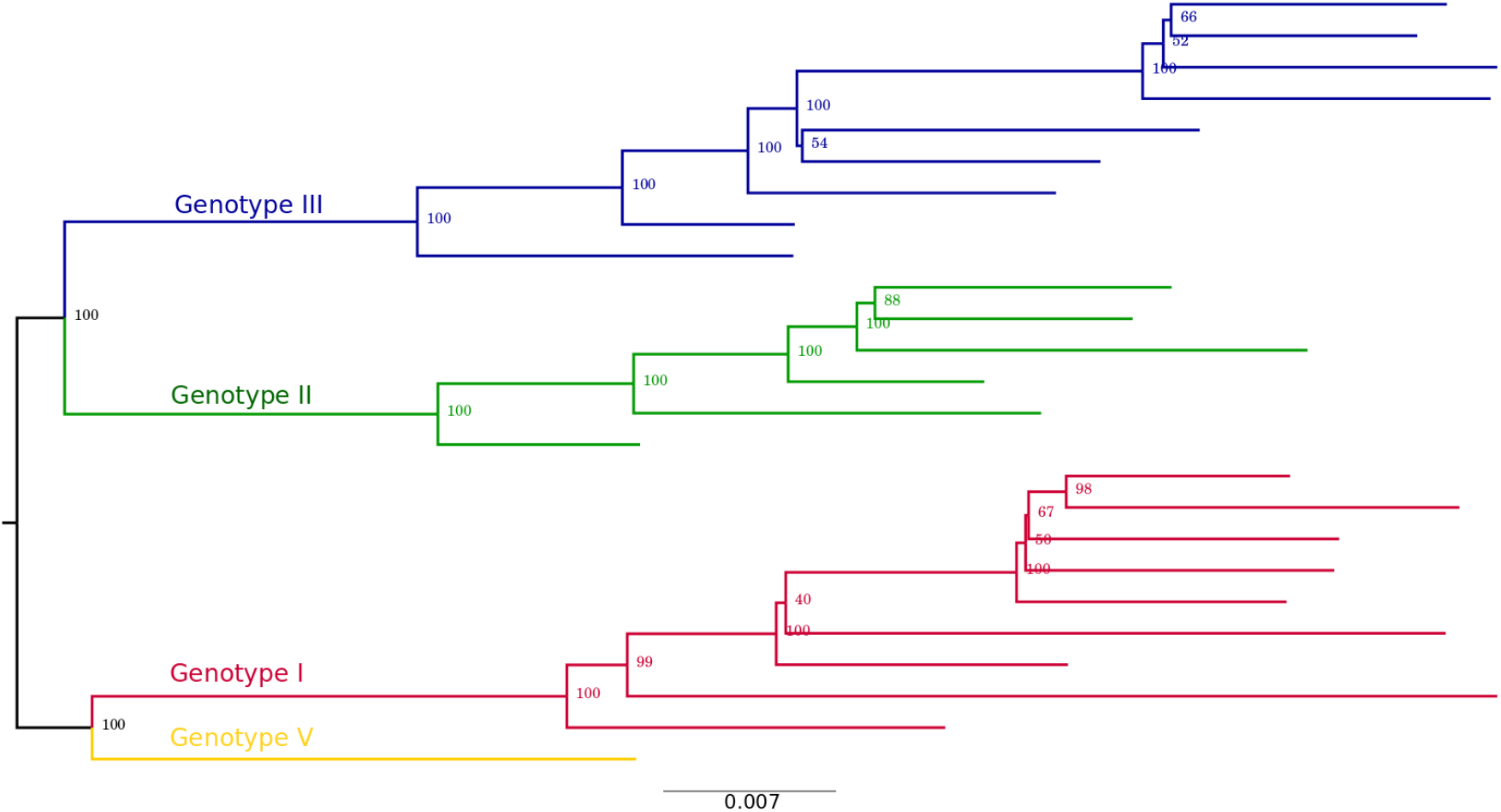
Maximum Likelihood inference of the multiple sequence alignment of the 25 DENV-3 complete genomes in the typing dataset. 807 complete DENV-3 genomes were clustered at 98% nucleotide identity and the representative genomes were aligned with mafft. A maximum likelihood tree was infered with RAxML. The tree is coloured according to genotype (red: genotype I; green: genotype II; blue: genotype III; purple: genotype IV).

**Figure S5.**
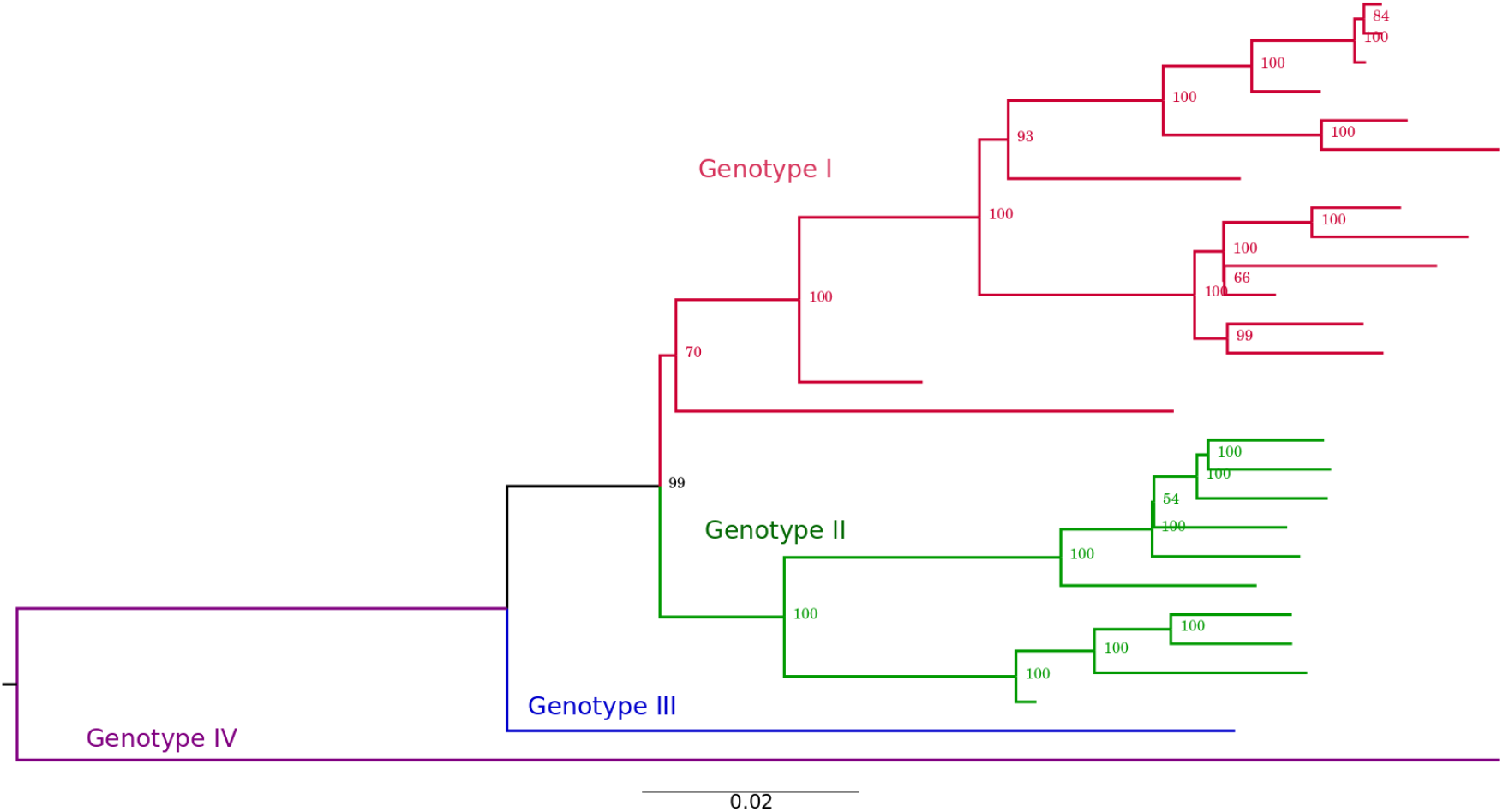
Maximum Likelihood inference of the multiple sequence alignment of the 27 DENV-4 complete genomes in the typing dataset. 320 complete DENV-4 genomes were clustered at 98% nucleotide identity and the representative genomes were aligned with mafft. A maximum likelihood tree was infered with RAxML. The tree is coloured according to genotype (red: genotype I; green: genotype II; blue: genotype III; purple: genotype IV).

