## Supplementary material for "DEN-IM: Dengue Virus identification from shotgun and targeted metagenomics": Table S1

**Table S1.** Overall alignment rate, in percentage, for the mapping against the DENV database, number of ORFs recovered, and respective serotype and genotype for 106 targeted sequencing samples.

| Sample | Raw MegaBases | % DENV DNA | CDS Assembly | Serotype | Genotype |
|---|---|---|---|---|---|
| SRR5821157 | 439.35 | 82.56 | consensus | 3 | III |
| SRR5821158 | 77.34 | 85.19 | consensus | 3 | III |
| SRR5821159 | 68.00 | 91.11 | consensus | 3 | III |
| SRR5821160 | 119.54 | 97.77 | consensus | 3 | III |
| SRR5821161 | 53.40 | 92.76 | consensus | 3 | III |
| SRR5821162 | 49.59 | 99.39 | consensus | 3 | III |
| SRR5821163 | 66.43 | 97.78 | consensus | 3 | III |
| SRR5821164 | 69.96 | 99.18 | consensus | 3 | III |
| SRR5821165 | 75.48 | 98.38 | consensus | 3 | III |
| SRR5821166 | 38.99 | 62.03 | *de novo* | 3 | III |
| SRR5821167 | 73.15 | 49.19 | *de novo* | 3 | III |
| SRR5821168 | 49.59 | 99.63 | consensus | 3 | III |
| SRR5821169 | 119.39 | 99.74 | *de novo* | 3 | III |
| SRR5821170 | 61.45 | 99.09 | consensus | 3 | III |
| SRR5821171 | 61.63 | 98.92 | consensus | 3 | III |
| SRR5821172 | 69.86 | 98.96 | *de novo* | 3 | III |
| SRR5821173 | 80.37 | 97.59 | *de novo* | 3 | III |
| SRR5821174 | 37.58 | 76.69 | *de novo* | 3 | III |
| SRR5821175 | 112.70 | 75.55 | *de novo* | 3 | III |
| SRR5821176 | 139.34 | 99.03 | *de novo* | 3 | III |
| SRR5821177 | 41.19 | 44.56 | *de novo* | 3 | III |
| SRR5821178 | 59.03 | 81.06 | *de novo* | 3 | III |
| SRR5821179 | 95.59 | 84.7 | *de novo* | 3 | III |
| SRR5821180 | 48.75 | 98.15 | consensus | 3 | III |
| SRR5821181 | 64.45 | 99.3 | consensus | 3 | III |
| SRR5821182 | 64.40 | 98.88 | consensus | 3 | III |
| SRR5821183 | 115.14 | 95.61 | consensus | 3 | III |
| SRR5821184 | 170.72 | 94.11 | *de novo* | 3 | III |
| SRR5821185 | 181.75 | 98.19 | *de novo* | 3 | III |
| SRR5821186 | 246.98 | 96.4 | *de novo* | 3 | III |
| SRR5821187 | 55.62 | 99.74 | consensus | 3 | III |
| SRR5821188 | 70.95 | 99.39 | consensus | 3 | III |
| SRR5821189 | 82.61 | 99.27 | *de novo* | 3 | III |
| SRR5821190 | 138.58 | 98.81 | consensus | 3 | III |
| SRR5821191 | 59.92 | 99.72 | *de novo* | 3 | III |
| SRR5821192 | 40.53 | 36.88 | consensus | 3 | III |
| SRR5821193 | 92.08 | 98.9 | *de novo* | 3 | III |
| SRR5821194 | 58.69 | 98.53 | consensus | 3 | III |
| SRR5821195 | 127.80 | 99.64 | consensus | 3 | III |
| SRR5821196 | 59.30 | 86.62 | *de novo* | 3 | III |
| SRR5821197 | 87.78 | 99.47 | *de novo* | 3 | III |
| SRR5821198 | 185.55 | 99.72 | *de novo* | 3 | III |
| SRR5821199 | 83.55 | 99.62 | consensus | 3 | III |
| SRR5821200 | 85.52 | 99.5 | consensus | 3 | III |
| SRR5821201 | 129.77 | 94.6 | consensus | 3 | III |
| SRR5821202 | 56.60 | 99.81 | consensus | 3 | III |
| SRR5821203 | 80.28 | 99.22 | consensus | 3 | III |
| SRR5821204 | 68.46 | 95.52 | *de novo* | 3 | III |
| SRR5821205 | 44.45 | 98.53 | consensus | 3 | III |
| SRR5821206 | 43.67 | 97.88 | consensus | 3 | III |
| SRR5821207 | 78.93 | 99.22 | *de novo* | 3 | III |

Continued on next page

**Table S1 – continued from previous page**

| Sample | Raw MegaBases | % DENV DNA | CDS Assembly | Serotype | Genotype |
|---|---|---|---|---|---|
| SRR5821208 | 87.45 | 97.72 | consensus | 3 | III |
| SRR5821209 | 73.40 | 94.16 | *de novo* | 3 | III |
| SRR5821210 | 55.86 | 91.35 | *de novo* | 3 | III |
| SRR5821211 | 75.53 | 85.6 | consensus | 3 | III |
| SRR5821212 | 98.89 | 99.09 | *de novo* | 3 | III |
| SRR5821213 | 84.85 | 95.03 | *de novo* | 3 | III |
| SRR5821214 | 15.33 | 96.28 | *de novo* | 3 | III |
| SRR5821215 | 13.08 | 96.74 | consensus | 3 | III |
| SRR5821216 | 45.07 | 98.85 | *de novo* | 3 | III |
| SRR5821217 | 161.65 | 88.94 | consensus | 3 | III |
| SRR5821218 | 51.09 | 95.29 | consensus | 3 | III |
| SRR5821219 | 84.68 | 99.1 | *de novo* | 3 | III |
| SRR5821220 | 88.26 | 82.64 | *de novo* | 3 | III |
| SRR5821221 | 64.76 | 86.62 | *de novo* | 3 | III |
| SRR5821222 | 93.47 | 97.48 | consensus | 3 | III |
| SRR5821223 | 86.50 | 98.99 | *de novo* | 3 | III |
| SRR5821224 | 73.31 | 26.43 | consensus | 3 | III |
| SRR5821225 | 68.85 | 98.43 | consensus | 3 | III |
| SRR5821226 | 67.75 | 96.67 | consensus | 3 | III |
| SRR5821227 | 32.56 | 99.54 | *de novo* | 3 | III |
| SRR5821228 | 38.73 | 86.68 | consensus | 3 | III |
| SRR5821229 | 77.18 | 99.69 | consensus | 3 | III |
| SRR5821230 | 175.73 | 99.58 | *de novo* | 3 | III |
| SRR5821231 | 100.82 | 99.58 | *de novo* | 3 | III |
| SRR5821232 | 86.89 | 99.47 | consensus | 3 | III |
| SRR5821233 | 270.15 | 99.56 | consensus | 3 | III |
| SRR5821234 | 76.07 | 99.75 | consensus | 3 | III |
| SRR5821235 | 32.78 | 79.78 | consensus | 3 | III |
| SRR5821236 | 80.19 | 24.72 | *de novo* | 3 | III |
| SRR5821237 | 50.59 | 97.38 | consensus | 3 | III |
| SRR5821238 | 63.56 | 97.63 | *de novo* | 3 | III |
| SRR5821239 | 29.66 | 41.15 | consensus | 3 | III |
| SRR5821240 | 62.61 | 94.64 | *de novo* | 3 | III |
| SRR5821241 | 17.52 | 98.03 | consensus | 3 | III |
| SRR5821242 | 58.86 | 99.25 | consensus | 3 | III |
| SRR5821243 | 50.08 | 93.56 | consensus | 3 | III |
| SRR5821244 | 32.67 | 99.09 | consensus | 3 | III |
| SRR5821245 | 64.96 | 99.77 | consensus | 3 | III |
| SRR5821246 | 104.11 | 90.14 | consensus | 3 | III |
| SRR5821247 | 98.64 | 99.73 | consensus | 3 | III |
| SRR5821248 | 129.28 | 90.73 | consensus | 3 | III |
| SRR5821249 | 45.76 | 93.13 | *de novo* | 3 | III |
| SRR5821250 | 72.54 | 98.88 | *de novo* | 3 | III |
| SRR5821251 | 115.85 | 97.7 | consensus | 3 | III |
| SRR5821252 | 60.76 | 94 | consensus | 3 | III |
| SRR5821253 | 64.45 | 99.66 | consensus | 3 | III |
| SRR5821254 | 0.27 | 98.12 | consensus | 3 | III |
| SRR5821255 | 62.53 | 99.55 | *de novo* | 3 | III |
| SRR5821256 | 54.57 | 99.58 | consensus | 3 | III |
| SRR5821257 | 34.90 | 99.53 | *de novo* | 3 | III |
| SRR5821258 | 68.64 | 99.6 | consensus | 3 | III |
| SRR5821259 | 73.04 | 98.8 | consensus | 3 | III |

Continued on next page

**Table S1 – continued from previous page**

| Sample | Raw MegaBases | % DENV DNA | CDS Assembly | Serotype | Genotype |
|---|---|---|---|---|---|
| SRR5821260 | 54.60 | 99.14 | consensus | 3 | III |
| SRR5821261 | 55.54 | 95.5 | *de novo* | 3 | III |
| SRR5821262 | 106.05 | 91.78 | consensus | 3 | III |
