## Supplementary material for "DEN-IM: Dengue Virus identification from shotgun and targeted metagenomics": Table S2

**Table S2.** Taxonomic profiling results for the target metagenomic samples with less than 70% DENV DNA.

| Sample | Bowtie2 | Kraken2 (minikraken2_V2 DB) | | |
|---|---|---|---|---|
| | **DENV (%)** | **Unclassified (%)** | ***Homo sapiens* (%)** | **DENV (%)** |
| SRR5821236 | 24.72 | 5.47 | 71.61 | 19.63 |
| SRR5821224 | 26.43 | 7.01 | 71.06 | 19.58 |
| SRR5821192 | 36.88 | 8.12 | 61.78 | 28.73 |
| SRR5821239 | 41.15 | 8.29 | 56.43 | 33.84 |
| SRR5821167 | 49.19 | 14.79 | 50.16 | 34.38 |
| SRR5821166 | 62.03 | 13.72 | 37.77 | 47.97 |
