## Supplementary material for "DEN-IM: Dengue Virus identification from shotgun and targeted metagenomics": Table S3

**Table S3.** Collection date, serotype confirmation and run accession identifier for the metagenomic sequecing dataset.

| Sample | Collection Date | Source | Serotype (qPCR) | Serotype | Genotype | Run Accession |
|---|---|---|---|---|---|---|
| 91-0104 | 21/9/2015 | plasma | 2 | 2 | III(AsianAmerican) | SRR8842525 |
| 91-0105 | 22/9/2015 | plasma | 2 | 2 | III(AsianAmerican) | SRR7252349 |
| 91-0115 | 30/9/2015 | plasma | 3 | - | - | SRR7252368 |
| 91-0118 | 5/10/2015 | plasma | 3 | 3 | III | SRR7252362 |
| 91-0132 | 19/10/2015 | plasma | 1 | 1 | V | SRR8883926 |
| 91-0135 | 27/10/2015 | plasma | 2 | 2 | III(AsianAmerican) | SRR9004764 |
| 92-1001 | 2/10/2015 | plasma | 1 | - | - | SRR7252337 |
| 92-1094 | 16/10/2015 | plasma | 2 | 2 | III(AsianAmerican) | SRR8842524 |
| CC0007 | 31/8/2010 | serum | 2 | 2 | III(AsianAmerican) | SRR7252354 |
| CC0009 | 31/8/2010 | serum | 3 | 3 | III | SRR8842527 |
| CC0010 | 27/8/2010 | serum | 3 | 3 | III | SRR7252358 |
| CC0011 | 27/8/2010 | serum | 3 | 3 | III | SRR8842526 |
| CC0030a | 1/9/2010 | serum | 4 | 4 | II | SRR7252356 |
| CC0030b | 1/9/2010 | serum | 4 | 4 | II | SRR7252355 |
| CC0031 | 2/9/2010 | serum | 2 | 2 | III(AsianAmerican) | SRR8842521 |
| CC0061 | 20/1/2011 | serum | 4 | 4 | II | SRR8842520 |
| CC0066 | 11/10/2011 | serum | 4 | 4 | II | SRR8842523 |
| CC0067 | 18/10/2011 | serum | 4 | 4 | II | SRR8842522 |
| CC0116 | 29/3/2012 | serum | 4 | 4 | II | SRR8842519 |
| CC0150 | 9/5/2012 | serum | 2 | 2 | III(AsianAmerican) | SRR8842518 |
| CC0186 | 17/7/2012 | serum | 4 | 4 | II | SRR9004763 |
| UCUG0186 | 30/8/2010 | serum | 4 | 4 | II | SRR8842528 |
| Negative Control | - | - | - | - | - | SRR8842530 |
| Positive Control | - | - | 2 | 2 | V(AsianI) | SRR8886136 |
| Spiked sample | - | - | 1,2,3,4 | 1,2,3,4 | V,III(AsianAmerican),III,II | SRR8842529 |
