## Supplementary material for "DEN-IM: Dengue Virus identification from shotgun and targeted metagenomics": Table S4

**Table S4.** DENV-1 Typing database

| Sample | ViPR Classification | Origin | Collection Year |
|---|---|---|---|
| EU482591 | DENV-1 V | USA | 2006 |
| KU509254 | DENV-1 V | Venezuela | 2011 |
| MF004384 | DENV-1 V | France | 2014 |
| GU131956 | DENV-1 V | Mexico | 2006 |
| AF311956 | DENV-1 V | Brazil | 1997 |
| FJ205874 | DENV-1 V | USA | 1995 |
| FJ478457 | DENV-1 V | USA | 1996 |
| EU482567 | DENV-1 V | USA | 1998 |
| DQ285559 | DENV-1 V | Reunion | 2004 |
| JN903578 | DENV-1 V | India | 2007 |
| KP188548 | DENV-1 V | Brazil | 2013 |
| JQ922544 | DENV-1 V | India | 1963 |
| KX380796 | DENV-1 V | Singapore | 2012 |
| JQ922548 | DENV-1 V | India | 2005 |
| KP406801 | DENV-1 V | South Korea | 2004 |
| DQ285562 | DENV-1 V | Comoros | 1993 |
| JQ922546 | DENV-1 V | India | 1971 |
| EF457905 | DENV-1 III | Malaysia | 1972 |
| AF180818 | DENV-1 II | Unknown | Unknown |
| JQ922547 | DENV-1 II | Thailand | 1960 |
| KY496855 | DENV-1 IV | Taiwan | 2016 |
| LC128301 | DENV-1 IV | Philippines | 2016 |
| KX951689 | DENV-1 IV | Taiwan | 2004 |
| KC762653 | DENV-1 IV | Indonesia | 2008 |
| KU509261 | DENV-1 IV | Indonesia | 2010 |
| AB189121 | DENV-1 IV | Indonesia | 1998 |
| KC762620 | DENV-1 IV | Indonesia | 2007 |
| EU863650 | DENV-1 IV | Chile | 2002 |
| AB195673 | DENV-1 IV | Japan | 2003 |
| AB204803 | DENV-1 IV | Japan | 2004 |
| JF459993 | DENV-1 I | Myanmar | 2002 |
| KT827371 | DENV-1 I | China | 2014 |
| KX620454 | DENV-1 I | China | 2014 |
| FJ639670 | DENV-1 I | Cambodia | 2001 |
| KU509250 | DENV-1 I | Thailand | 2012 |
| KJ755855 | DENV-1 I | India | 2013 |
| GU131678 | DENV-1 I | Viet Nam | 2008 |
| KU509265 | DENV-1 I | Unknown | 2012 |
| KF955446 | DENV-1 I | Viet Nam | 2008 |
| JF937615 | DENV-1 I | Viet Nam | 2008 |
| FJ639678 | DENV-1 I | Cambodia | 2003 |
| EU660395 | DENV-1 I | Viet Nam | 2007 |
| AB608789 | DENV-1 I | Taiwan | 1994 |
| GQ868636 | DENV-1 I | Cambodia | 2008 |
| KY586539 | DENV-1 I | Thailand | 1995 |
| KU509258 | DENV-1 I | Eritrea | 2010 |
