## Supplementary material for "DEN-IM: Dengue Virus identification from shotgun and targeted metagenomics": Table S5

**Table S5.** DENV-2 Typing database

| Sample | ViPR Classification | Origin | Collection Year |
|---|---|---|---|
| HQ705624 | DENV-2 III (AsianAmerican) | Nicaragua | 2009 |
| KY977454 | DENV-2 III (AsianAmerican) | Panama | 2011 |
| KY474330 | DENV-2 III (AsianAmerican) | Ecuador | 2014 |
| FJ024473 | DENV-2 III (AsianAmerican) | Colombia | 2005 |
| JX669476 | DENV-2 III (AsianAmerican) | Brazil | 2010 |
| JN819419 | DENV-2 III (AsianAmerican) | Brazil | 2000 |
| KF955364 | DENV-2 III (AsianAmerican) | Puerto Rico | 2006 |
| JX669480 | DENV-2 III (AsianAmerican) | Brazil | 1995 |
| FJ639699 | DENV-2 III (AsianAmerican) | Cambodia | 2002 |
| EU482449 | DENV-2 III (AsianAmerican) | Viet Nam | 2006 |
| EU482778 | DENV-2 III (AsianAmerican) | Viet Nam | 2003 |
| KY586692 | DENV-2 V (AsianI) | Thailand | 2001 |
| KY586679 | DENV-2 V (AsianI) | Thailand | 2001 |
| KY586571 | DENV-2 V (AsianI) | Thailand | 2006 |
| KY586572 | DENV-2 V (AsianI) | Thailand | 2006 |
| EU726767 | DENV-2 V (AsianI) | Thailand | 1994 |
| GQ868591 | DENV-2 V (AsianI) | Thailand | 1964 |
| KF704356 | DENV-2 IV (AsianII) | Cuba | 1981 |
| JQ922552 | DENV-2 I (American) | India | 1960 |
| KJ918750 | DENV-2 I (American) | India | 2007 |
| JQ922553 | DENV-2 I (American) | India | 1980 |
| GQ868592 | DENV-2 I (American) | Colombia | 1986 |
| JX966379 | DENV-2 I (American) | Mexico | 1994 |
| GQ398257 | DENV-2 I (American) | Indonesia | 1977 |
| KY923048 | DENV-2 VI (Sylvatic) | Malaysia | 2015 |
| JF260983 | DENV-2 VI (Sylvatic) | Spain | 2009 |
| KY937189 | DENV-2 II (Cosmopolitan) | China | 2015 |
| KY937188 | DENV-2 II (Cosmopolitan) | China | 2015 |
| KY937187 | DENV-2 II (Cosmopolitan) | China | 2015 |
| JQ955624 | DENV-2 II (Cosmopolitan) | India | 2011 |
| KU509271 | DENV-2 II (Cosmopolitan) | India | 2006 |
| KF041232 | DENV-2 II (Cosmopolitan) | Pakistan | 2011 |
| JQ922551 | DENV-2 II (Cosmopolitan) | India | 2005 |
| JX475906 | DENV-2 II (Cosmopolitan) | India | 2009 |
| MG779194 | DENV-2 II (Cosmopolitan) | Kenya | 2017 |
| FJ882602 | DENV-2 II (Cosmopolitan) | Sri Lanka | 1996 |
| EU056810 | DENV-2 II (Cosmopolitan) | Burkina Faso | 1983 |
| KY627763 | DENV-2 II (Cosmopolitan) | Burkina Faso | 2016 |
| KM279515 | DENV-2 II (Cosmopolitan) | Singapore | 2011 |
| KX452015 | DENV-2 II (Cosmopolitan) | Malaysia | 2014 |
| KC762662 | DENV-2 II (Cosmopolitan) | Indonesia | 2007 |
| KU509270 | DENV-2 II (Cosmopolitan) | Unknown | 2012 |
| KP012546 | DENV-2 II (Cosmopolitan) | China | 2014 |
| KX452034 | DENV-2 II (Cosmopolitan) | Malaysia | 2014 |
| KX452048 | DENV-2 II (Cosmopolitan) | Malaysia | 2014 |
| KX452044 | DENV-2 II (Cosmopolitan) | Malaysia | 2014 |
| HM488257 | DENV-2 II (Cosmopolitan) | Guam | 2001 |
| KU509277 | DENV-2 II (Cosmopolitan) | Philippines | 2010 |
| KU509269 | DENV-2 II (Cosmopolitan) | Philippines | 2009 |
| KU509274 | DENV-2 II (Cosmopolitan) | Philippines | 2010 |
| GQ398263 | DENV-2 II (Cosmopolitan) | Indonesia | 1975 |
