## Supplementary material for "DEN-IM: Dengue Virus identification from shotgun and targeted metagenomics": Table S6

**Table S6.** DENV-3 Typing database

| Sample | ViPR Classification | Origin | Collection Year |
|---|---|---|---|
| KF954946 | DENV-3-III | China | 2013 |
| JQ922557 | DENV-3 III | India | 2005 |
| KU509286 | DENV-3 III | India | 2011 |
| EU687233 | DENV-3 III | USA | 2002 |
| GQ252674 | DENV-3 III | Sri Lanka | 1997 |
| FJ882573 | DENV-3 III | Sri Lanka | 1993 |
| GQ199887 | DENV-3 III | Sri Lanka | 1983 |
| JQ922555 | DENV-3 III | India | 1966 |
| HM631854 | DENV-3 II | Cambodia | 2008 |
| KY586703 | DENV-3 II | Thailand | 2006 |
| KU509280 | DENV-3 II | Thailand | 2011 |
| FJ744730 | DENV-3 II | Thailand | 2001 |
| KY586814 | DENV-3 II | Thailand | 2006 |
| DQ863638 | DENV-3 II | Thailand | 1973 |
| KC762684 | DENV-3 I | Indonesia | 2007 |
| KY863456 | DENV-3 I | Indonesia | 2016 |
| KC762691 | DENV-3 I | Indonesia | 2008 |
| KC762692 | DENV-3 I | Indonesia | 2010 |
| KY794787 | DENV-3 I | Papua New Guinea | 2007 |
| MF004386 | DENV-3 I | Malaysia | 2012 |
| AB189128 | DENV-3 I | Indonesia | 1998 |
| KU509279 | DENV-3 I | Philippines | 2008 |
| FJ898455 | DENV-3 I | Cook Islands | 1991 |
| KU725666 | DENV-3 V | Unkown | Unknown |
