## Supplementary material for "DEN-IM: Dengue Virus identification from shotgun and targeted metagenomics": Table S7

**Table S7.** DENV-4 Typing database

| Sample | ViPR Classification | Origin | Collection Year |
|---|---|---|---|
| MG601754 | DENV-4 I | China | 2013 |
| KY586839 | DENV-4 I | Thailand | 1995 |
| KT026308 | DENV-4 I | Thailand | 2011 |
| JN638572 | DENV-4 I | Cambodia | 2008 |
| KY586942 | DENV-4 I | Thailand | 2006 |
| KP792537 | DENV-4 I | Singapore | 2011 |
| MG272273 | DENV-4 I | India | 2016 |
| MG272272 | DENV-4 I | India | 2016 |
| KU509287 | DENV-4 I | India | 2009 |
| JQ922559 | DENV-4 I | India | 1979 |
| GQ868594 | DENV-4 I | Philippines | 1956 |
| JQ922558 | DENV-4 I | India | 1962 |
| KU523872 | DENV-4 II | Indonesia | 2015 |
| KP723482 | DENV-4 II | China | 2010 |
| JX024757 | DENV-4 II | Singapore | 2010 |
| KC762695 | DENV-4 II | Indonesia | 2007 |
| JQ915088 | DENV-4 II | New Caledonia | 2009 |
| GQ398256 | DENV-4 II | Singapore | 2005 |
| KP188557 | DENV-4 II | Brazil | 2012 |
| KY474335 | DENV-4 II | Ecuador | 2014 |
| KT276273 | DENV-4 II | Haiti | 2014 |
| KF907503 | DENV-4 II | Senegal | 1953 |
| KY586945 | DENV-4 III | Thailand | 1998 |
| JF262779 | DENV-4 IV | Malaysia | 1975 |
